# MLS3: A New Type of Multilevel Selection

**DOI:** 10.1101/2020.10.22.350272

**Authors:** Christopher T. Jones, W. Ford Doolittle, Edward Susko, Letitia Meynell, Celso A. Neto, Timothy M. Lenton, Joseph P. Bielawski

## Abstract

In most multispecies multilevel selection (MLS) models, offspring communities are generated by random assembly of individuals in numbers reflecting sizes of parental communities releasing them (MLS1), or by differential community dispersal based on a community-level trait such as size (MLS2). In both, offspring communities *colonize* vacant spaces: different communities never compete for the same space. Here we propose a third MLS type (MLS3) where multispecies communities disperse (*migrate*) into already-occupied spaces, larger communities more frequently. Conspecific variants compete, often opposing selection for community size against fitness within species. This makes the outcome of MLS3 less apparent than MLS1 and MLS2 where such tension is absent. Our simulations show that, if community size depends strongly on reduction in the fitness of individual community members, such a reduction (comprising a sort of “inter-species altruism”) will evolve. The framework we present represents a step toward conceptualizing *community coalescence* in the context of metacommunities.

## Introduction

It has long been recognized that “natural selection can operate at more than one level of the biological hierarchy, each level favoring a different set of traits” (1). However, even if “traits can evolve by virtue of benefiting whole groups, despite being selectively disadvantageous within groups” (1), they are unlikely to do so without the imposition of some mechanism that can overpower the selfish tendencies of within-group natural selection (the tragedy of the commons, 2). Models of multilevel selection (MLS) that bestow the conditions required for differential community-level reproduction have been proposed. Many are based on some kind of “ecological scaffolding”, defined as a set of assumptions or procedures (e.g., features of the ecological milieu) that “exogenously impose Darwinian-like properties onto communities” (3,4). Following “Lewontin’s Recipe” (5), they require “that variation among communities is discrete, that communities replicate and that offspring communities show some resemblance to parental communities” (6).

In this paper we use simulations to explore possible mechanisms of multilevel selection in the context of a network of ecological patches that can each support a multi-species microbial community (i.e., together a metacommunity, 7,8,9). Existing models of MLS (5), as commonly brought to bear on the evolution of altruism within species (10,11), can generally be categorised as either MLS1 or MLS2 (12,13). MLS1 is arguably not selection of groups but rather selection of individuals or their genes by virtue of group membership (13). In our multi-species context, MLS1 might be said to apply when new communities are formed in vacant patches by random assembly of species members (individual organisms) drawn from all (or all adjacent) extant “parental” communities. For selection to occur, it must be that some communities, because of their species composition, produce more individual organisms than others. MLS2, on the other hand, requires some communities to be more likely to form new colonies via the dispersal of some portion of their cells into vacant patches. Colonization is thus a form of “reproduction” at the community level, conforming to Lewontin’s Recipe. In our metacommunity context in which the number of patches is finite, space for colonization by random assemblies drawn from multiple parental communities (MLS1) or a single parental community (MLS2) is provided by episodic culling of existing communities. Under this condition, both MLS1 and MLS2 support an increase in the fraction of communities whose species composition corresponds to larger constituent populations (MLS1) or a greater likelihood of community dispersal (MLS2).

A third mechanism of MLS (MLS3) that is distinct from existing models incorporates the following conditions. (i) That new communities are formed when a portion of a donor community is transferred into a neighboring recipient patch that is already occupied (i.e., via *migration* instead of *colonization*). (ii) That the probability that a species with a particular genotype from the donor community displaces its competitor in the recipient patch (a member of the same species but with a different genotype) is a function of both the size of its population in the donor community and the nature of its competitor in the recipient community (the difference in fitness of the two genotypes). These assumptions, which are likely to be consistent with what can occur in natural systems (e.g., by “community coalescence”, 14), potentially place selection within an individual species in direct conflict with selection at the level of the community and makes the outcome less obvious compared to standard models of MLS. Nevertheless, we show via simulation that (i) and (ii) can lead to an increase in the size of the constituent populations of a community at the expense of the fitness of individual species even when as many as 20 species are required for successful community function.

## Background

Imagine a single-species population composed of two genetic variants, which to align with much of the literature we call a selfish type and an altruistic type. The selfish type is individually fitter (produces more offspring) in comparison to the altruistic type in any group, but a group with more altruists will initially grow faster (acquire more members) than a group with fewer. However, any single mixed group will eventually be reduced to nothing but the selfish type because of differential reproduction of types. In other words, “the inherent logic of the commons remorselessly generates tragedy” (2) due to “subversion from within” (15). To explain the appearance of altruism in nature, Hamilton proposed his theory of inclusive fitness (16,17), a key feature of which is that it can promote altruism within a single population (albeit structured by kin groups). An alternative, known as trait-group selection (10), is aligned with MLS1. This envisions that some groups in a population of groups produce more individuals than others by virtue of their composition, namely if they include many altruists. Thus, the next generation of groups, produced by random recruitment from a pool of all individuals released from the previous generation, will contain higher proportions of altruists. Sober (11) proposed a third mechanism based on the assumptions that groups can reproduce as groups by colonizing new space (i.e., by MLS2), and that larger groups tend to be more successful colonizers. Any of these alternatives can support an increase in the mean proportion of altruists in a population over time.

Here we aim to extend the logic of MLS to metacommunities with migration between patches. Our model is based on communities composed of multiple species of microorganisms whose combined metabolic processes are assumed to form a mutualistic network of chemical transformations of unspecified topology (e.g., a closed nutrient-recycling loop). The microbial perspective was chosen in part to facilitate a comparison between our metacommunity approach and the numerous models that explore the evolution of single microbial communities (18,19,20,21). We assume that multiple variants exist within each species, differing in fitness as determined by alleles that impact their reproduction rates (e.g., by varying their consumption and growth rates). Differences between variants provide the bases for selection to act within species and would, if there were no community-level selection, always favour the fixation of alleles conferring higher consumption and growth. We also assume that the size of a community (and consequentially, the size of its migration propagules that impact the probability of successful dispersal) is a function of its degree of self-regulation. That is, we assume that community size can be maximized only when the consumption and growth rates of its constituent species achieve a particular balance or “target” configuration, most easily imagined to be that which leads to no excessive accumulation of any intermediate substrate or that maximizes throughput. It is easy to further imagine that this balance can be achieved only at the expense of the fitness of individuals in one or more constituent species, each of which plays a role similar to that of altruists in the single species scenario (cf., 10,11). The suppression of fitness is consistent with the biologically reasonable assumption that maximum consumption and growth rates differ between species and cannot always be sustained in a community setting for specific biochemical, physiological, or environmental reasons. These assumptions force a trade-off between what is best for an individual within a species (maximizing its fitness) and what is best for the community (maximizing its ability to disperse as a community).

Community structure is imposed in our model, which does not account for how nutrient recycling might arise in the first place. We also avoid complexities associated with intracommunity interactions under scenarios more complex than mutualistic nutrient recycling. Simplifying assumptions of this kind, in which within-community dynamics are not considered in detail but between-community dynamics are extensively considered, are common in discussions of MLS (e.g., the “patch-dynamic” paradigm, 8). An alternative approach would be to specify detailed mechanisms of species-specific consumption, growth, reproduction and death, interactions between species and interactions between a community and its abiotic environment. Mechanistic agent-based models of this kind make within-community dynamics explicit and have been used to explore the way nutrient recycling might arise de novo and how a community might evolve to constrain elements of the abiotic environment to maintain community viability (18,19,20,21). Our objective is to demonstrate that MLS3 can *in principle* (i.e., under idealized conditions) impose a selection regime that favors the assembly of larger communities even when this comes at the expense of the fitness of one or more community members in a manner consistent with Wilson (1975) and Sober (1988). We argue that our approach is appropriate because the details of community-level dynamics can vary without changing MLS3 as a mechanism of community-level selection. A full description of our model is provided in Methods.

## Results

Simulations were conducted on a network of 400 ecological patches each of which supports a community of microbes. Each community was assumed be composed of variants of the same 20 species. Each variant was assigned a random fitness coefficient *F*_*i*_, and each community was therefore specified by a vector of fitnesses ***F*** = ⟨*F*_1_, …, *F*_20_⟩. The nominal effective size of each population in a community was assumed to be a function of the scaled distance between ***F*** and a vector of target fitnesses ***T***, the maximum size (1 × 10^6^) corresponding to ***F*** = ***T***. The increase in population size with each incremental decrease in the distance between ***F*** and ***T*** was a model parameter, as was the mean size of random variations applied to the population size at each timestep. Models for MLS1, MLS2 and MLS3 were applied to the same initial set of 400 randomly assembled communities. Each model induced a change in the initial cloud of 400 ***F*** vectors over time, corresponding to a change in the distribution of the nominal population size. Figure 1 shows such distributions as a function of the scaled distance between ***F*** and ***T*** at the start of each simulation and after *t* = 1000 timesteps (*t* = 2000 in Figure 1F). Each panel represents simulations conducted under different parameter settings and assumptions as listed in Table 1. See Methods for details.

**Table 1:** Parameter settings for each batch of simulations. The distance between *T* and **1** is proportional to δ, larger values of which reduce the ability of MLS3 to oppose ENS at the level of individuals or alleles within species (see Figure 2). The size of each species population in a patch is increased or decreased randomly at each timestep by a mean proportion ρ. The addition of this random variation to population size reduces the negative correlation between community regulation and Dist(***T***, ***F***(*t*)) and concomitantly the ability of MLS3 to oppose ENS. The stringency of the relationship between Dist(***T***, ***F***(*t*)) and community size is controlled by σ, where larger values decrease the gain in community size with each incremental step of ***F*** toward ***T***. Making σ larger reduces the ability of MLS3 to oppose ENS. Two variants of MLS3 were implemented, one in which migration propagules contain cells from all 20 species (full), and one in which migration propagules contain a random selection of species (partial) drawn from the donor community.

| Simulation | $\delta$ (fitness) | $\rho$ (population size) | $\sigma$ (stringency) | MLS3 Migration |
| --- | --- | --- | --- | --- |
| 1 (figure 2A) | $1 \times 10^{-6}$ | 0.01 | 0.20 | full |
| 2 (2B) | $1 \times 10^{-6}$ | 0.10 | 0.20 | full |
| 3 (2C) | $5 \times 10^{-6}$ | 0.01 | 0.20 | full |
| 4 (2D) | $5 \times 10^{-6}$ | 0.10 | 0.20 | full |
| 5 (2E) | $1 \times 10^{-6}$ | 0.01 | 1.00 | full |
| 6 (2F) | $1 \times 10^{-6}$ | 0.01 | 0.20 | partial |

**Figure 1:**
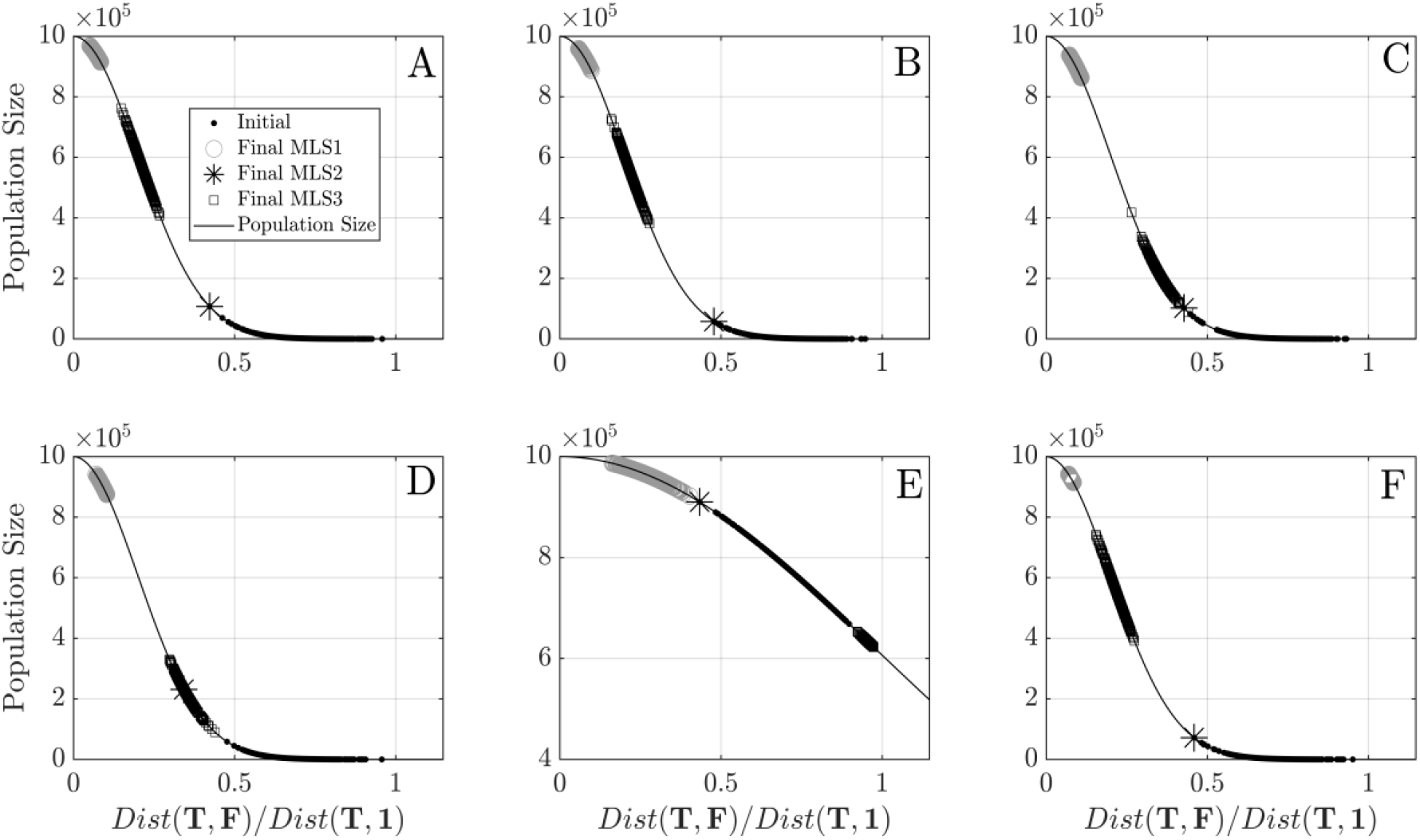
Each plot shows the distribution of the nominal effective population size for all patches as a function of the scaled distance at the start of each simulation (black dots) and at the end of each simulation under MLS1 (gray circles), MLS2 (black asterisk) and MLS3 (black squares). Parameter settings for each panel are listed in Table 1. A given method of MLS was effective in regulating community size in any case where the distribution of distance ratios *x*(*t*) = Dist(***T, F***(*t*))/Dist(***T***, **1**) was closer to zero by the end of the simulation than it was at the beginning of the simulation. See texts for a complete description of results. Additional figures are provided in Supplementary Material.

In Figure 1A the initial set of ***F*** vectors started some distance from the target ***T***. Under MLS1 all vectors were close to target by the end of the simulation, corresponding to distances close to zero and nominal population sizes close to 1 × 10^6^. Under MLS2 the system collapsed to the single ***F*** vector nearest ***T***, meaning that all 400 communities were identical. In this case the common population size was approximately 0.1 × 10^6^. Under MLS3 the ***F*** vectors moved toward ***T*** corresponding to a mean nominal population size of approximately 0.5 × 10^6^, but was less effective than MLS1. It is evident that the tension between ENS favoring greater individual fitness and MLS favoring larger populations prevented the ***F*** vectors from moving as close to ***T*** under MLS3 as they did under MLS1 where the tension was absent.

In Figure 1B simulations included larger random variations in population size. This was expected to make it more difficult for MLS3 to force the initial set of ***F*** vectors closer to ***T*** by reducing the correlation between distance and population size. However, the difference between Figure 1A and Figure 1B is negligible. In Figure 1C the reduction in fitness required to attain maximum population size was increased. This was also expected to make it more difficult for MLS3 to force the initial set of ***F*** vectors closer to ***T***. And indeed, although MLS1 and MLS2 appear to have been as effective as they were in Figure 1A and 1B in increasing the mean nominal population size, MLS3 was clearly less effective than before, attaining a mean size of approximately 0.2 × 10^6^, less than half of its previous value. Similar results are shown in Figure 1D.

Figures 1A to 1D demonstrate that MLS3 can provide an effective counter to ENS within species to promote an increase in population size. Its ability to do so, however, is grounded in the assumption that the probability that a donor species will displace its counterpart in the recipient patch is partly a function of the size of the donor community. This assumption is more effective when each incremental decrease in the distance between ***F*** and ***T*** corresponds to a larger increase in population size. In Figure 1E the increase in population size was reduced compared to previous simulations. Consequently, whereas MLS1 and MLS2 were as effective as in previous simulations, the mean nominal population size actually decreased under MLS3 in Figure 1E from an initial value of approximately 0.75 × 10^6^ to a value slightly more than 0.6 × 10^6^.

Our simulations of MLS3 were to this point conducted with the assumption that a migration propagule always includes cells from all 20 species in the donor community. This was meant to put MLS3 on an equal footing with MLS2 under which a donor community can always be replicated by colonizing an empty patch. Even with this assumption, however, a donor community is usually not replicated in a recipient patch under MLS3 because fixation is stochastic. Migration under MLS3 is therefore typically incomplete (i.e., only some migrants are fixed in the recipient patch). This suggests that MLS3 might be effective even under partial migration, where propagules contain only some fraction of the species in the donor community. To test this, the MLS3 algorithm was modified to make each migration propagule contain a random sample of the 20 species from the donor community (but each still at 10% of the population size). Results are shown in Figure 1F, where the parameter settings were the same as in Figure 1A but with partial migration. MLS3 was about as effective with partial migration as it was under full migration. However, it took twice as many times steps (i.e., 2000 instead of 1000) for the system to reach a stationary state with partial migration (see Figure 6 in Supplementary Material).

## Discussion

The “tragedy of the commons” (2) is rooted in the fact that what is advantageous to an individual is not necessarily advantageous to the group. In the context of a community of microbes, the tragedy might be expressed by the tendency of individual species to selfishly evolve to maximize their rates of reproduction even when larger populations might be achieved by all community members if only some species would hold this tendency in check. Multilevel selection provides a solution to the tragedy of the commons. However, standard formulations of MLS are typically based on the assumption that offspring communities are formed in vacant niches (i.e., by colonization). It is rather obvious that any community-level property that increases the probability of successful dispersal by colonization will tend to be selected, and that the probability of successful colonization will improve over time. This is because, without intraspecies competition as would occur under migration, offspring communities are more likely to resemble parental types. Dispersal of communities by colonization may well play an important role in the regulation of community-level properties (3) or in the evolution of individuality (4). However, in a mature biosphere in which living systems are ubiquitous it might often be the case that dispersal is by migration into locations that are already occupied, or what Rillig and colleagues call community coalescence (14). In such cases it is less apparent that MLS can act to overcome individual level selection for maximum fitness.

Dispersal by migration has been explored using mechanistic agent-based simulations. For example, in what might fairly be called the Network Flask Model (22) multiple microbial systems or flasks were linked in a circular topology and allowed to exchange a limited amount of material at each time step. Simulations suggest that such communication between flasks can improve nutrient recycling and temperature regulation (both being community-level properties) over time. However, substantial genetic drift caused by small populations and high mutation rates may have obscured the full impact of dispersal in that study. In a more recent study, microbial communities were assumed to be composed of a source species that converts substrate X to Y, a mutualistic species that converts Y back to X at some cost to fitness *κ*, and a parasitic species that converts Y to an unusable waste product Z (23). Community size could be increased by increasing the proportion of mutualists in the system due to their recycling of Y back to X. Nevertheless, any single community was typically dominated by the parasitic species when recycling was costly (*κ* > 0). The mutualists only dominated in a metacommunity setting in which neighboring flasks were permitted to exchange a fixed proportion of their contents (cf. 11). Results of this kind illustrate how migration can support selection for a community-level property. However, in the first example community self-regulation was not assumed to require a reduction in the fitness of some community members (i.e., there was no altruism). Instead, nutrient recycling and regulation emerged from the combined metabolic by-products of all species within a community. In the second example migration did not involve competition between different variants of the same species.

Under the model of MLS3 presented here, natural selection at the species level was explicitly assumed to oppose selection for community size. This was implemented by making community size a function of the difference between the vector ***F*** of fitnesses for the species within a community and a target vector ***T*** meant to represent an optimal balance between the consumption and growth rates of all constituent species. Community regulation consisted of attaining a vector ***F*** closer to ***T***, corresponding to a larger community size and a greater probability of dispersal. Since it was assumed that ***T*** < **1**, regulation could only be achieved at the expense of the fitness of individuals within species. Our simulations show that such regulation can be achieved under MLS3 when the change in community size is sensitive to changes in the difference between ***F*** and ***T*** (i.e., when σ = 0.20) but is ineffective when this is not the case (i.e., when σ = 1.00). An example of regulation of the kind considered here would be a community of microbes whose metabolic processes impact an environmental parameter (e.g., temperature, pH, chemical ratios, etc.) that has a knock-on effect on community size. In such a case it might be necessary to hold the consumption and growth rates of some community members in check to maintain viability. Our results suggest that, if the relationship between the size of a community (or more generally, its ability to disperse) is strongly dependent on the maintenance of such parameters within a narrow range of values (corresponding to a smaller σ in our model) then MLS3 can support community regulation. Our model is highly idealized, however, so it is not clear how effective MLS3 might be in real metacommunity systems. Nevertheless, our results show that the ecological milieu upon which MLS3 is based (i.e., where well-regulated communities with larger populations are more likely to donate propagules to neighboring patches) does not necessarily require the assumption of dispersal by colonization but can be effective even when dispersal is by migration.

A note on the definition of an ecological scaffolding is warranted. The term was initially defined to be any set of assumptions or procedures that bestow the conditions required for selection at the level of multicellular collectives, such as differential community-level reproduction (3, 4). The reference to ecology emphasizes the importance of ecological dynamics such as life history traits or mechanisms of dispersal in a metapopulation that are typically marginalized under traditional models (i.e., those that assume an idealized Wright-Fisher population, 24, 25). A set of procedures can be called an ecological scaffolding only if it can be shown to result in the emergence of a Darwinian population (26) in which communities act as Darwinian individuals (26). This is arguably not the case under the present model of MLS1 because it is individual species that disperse at differential rates, not the community, although the rate of dispersal is dependent on community size. It clearly is the case under the present model of MLS2, where communities are reproduced in vacant patches at differential rates that depend on community size, and where each offspring community exactly resembles its parent. It is also arguably the case under MLS3 with full migration because an offspring community is a combination of species drawn from two parental communities, and its size will therefore often be correlated with the size of at least one of its parents (e.g., the one that was selected for migration). The case is perhaps less clear under MLS3 with partial migration, which in some ways is not unlike MLS1. The point is that, although MLS can increase the number of well-regulated communities that each bear a close resemblance to one another in size, it is not necessary for communities to be Darwinian individuals for this to occur. An ecological scaffolding might therefore be more broadly construed as any set of assumptions or procedures that bestow the conditions required for differential community *reassembly* (not necessarily community reproduction). Under this definition MLS1 and MLS3 with partial migration both comprise an ecological scaffolding. Although community-level selection does not act in either case, well-regulated communities are gradually reassembled via a series of dispersal events.

## Methods

A community is assumed to consist of a collection of 20 different species whose mutualistic interactions are required to maintain community viability. Each species within a given community is assigned a fitness coefficient (e.g., *F*_*i*_ for the *i*^*th*^ species) that reflects its consumption and growth rates compared to conspecifics in other communities. These are held constant once assigned under the assumption that the timespan of each simulation is short enough that mutations that impact *F*_*i*_ are unlikely to occur. Two variants of the same species from different communities assigned fitnesses *F*_*i*_ and 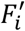 are assumed to differ in their consumption and growth rates as determined by differences in one or more unspecified alleles. The fitter variant is assumed to reproduce at a higher rate, all else being equal. However, the relationship between fitness and reproduction rate is unspecified. Instead, whenever two variants of the same species occupy the same patch, one or the other is assumed to eventually be fixed with a probability that depends on the difference 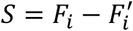 (i.e., the selection coefficient), as specified below. Fitness values are assumed to be in the interval 1 − δ ≤ *F*_*i*_ ≤ 1 where δ determines the difference between the fittest and least fit variant within any given species. The simulations reported here were conducted with δ ∈ {1 × 10^−6^, 5 × 10^−6^}. A community is identified by the vector ***F*** = ⟨*F*_1_, …, *F*_20_⟩ that specifies the fitnesses of each of its constituent species. The vector of maximum fitnesses is the ones vector **1** = ⟨1, …,1⟩.

All twenty species in any community are assumed to attain a fixed population size by the end of each model timestep, consistent with a state of dynamic equilibrium. For simplicity, this size is assumed to be the same for all species in the community, although our results do not depend on this assumption. Population size is assumed to be a function of the distance between the vector ***F*** assigned to a community and a vector ***T*** = ⟨*T*_1_, …, *T*_20_⟩ of target values:

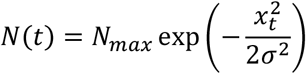

Here *x*_*t*_ is the Euclidean distance *Dist*(***T, F***(*t*)) between ***T*** and ***F***(*t*) divided by *Dist*(***T*, 1**) computed at timestep *t*, and *N*_*max*_ = 1 × 10^6^ is the maximum population size permitted under the model before random effects are applied. The nominal size of a community, which is 20 × *N*(*t*), increases as its assigned vector ***F*** moves closer to the target vector ***T*** and reaches its maximum value of 20 × *N*_*max*_ when ***F*** = ***T*** or *x*_*t*_ = 0. The parameter σ was selected to generate scenarios in which *N*(*t*) was either more (σ = 0.20) or less (σ = 1.00) sensitive to changes in *Dist*(***T, F***(*t*)), and can be thought of as a measure the stringency of selection against poorly-regulated communities, those with ***F*** further from ***T***. To add noise to the system, the size of each population in any given community was increased or decreased at each timestep by the same random proportion drawn from a beta distribution with expected value selected from *ρ* ∈ {0.01, 0.10}. The purpose was to generate scenarios in which *N*(*t*) was either more (*ρ* = 0.01) or less (*ρ* = 0.10) dependent on *Dist*(***T, F***(*t*)), the former being the case in which MLS3 is likely to be more effective. Note that this noise is multiplicative rather than additive.

The target ***T*** represents the specific community configuration (e.g., a balance between the consumption and growth rates of all constituent species) that maximizes community size and consequently the ability of a community to disperse. A well-regulated community is one for which ***F*** is closer to ***T***. If the target for each species was set to its maximum value (i.e., if *T*_*i*_ = 1, *i* ∈ {1, …,20} or ***T*** = ⟨1, …,1⟩ = **1**) then what is best for an individual within a species (i.e., maximizing its fitness) would be aligned with what is best for the community (i.e., maximizing its ability to disperse as a community). The resulting lack of conflict would be insufficient to test MLS3 since self-regulation within any single community would be possible via evolution by natural selection (ENS) acting within each species to maximize fitness. To create a tension between the individual and the community comparable to that in models for the evolution of altruism (e.g., 11), it must be assumed that ***T*** < **1** (i.e., some or all *T*_*i*_ < 1). Under this assumption, a community can increase its ability to reproduce as a community by attaining a larger community size only when it includes one or more species whose rate of reproduction is less than its species-specific maximum rate (i.e., with *F*_*i*_ < 1 for some or all *i* ∈ {1, …,20}). In this case regulation cannot be easily achieved without some form of MLS that can oppose ENS acting at the level of individuals or upon alleles within species, as illustrated in Figure 2.

**Figure 2:**
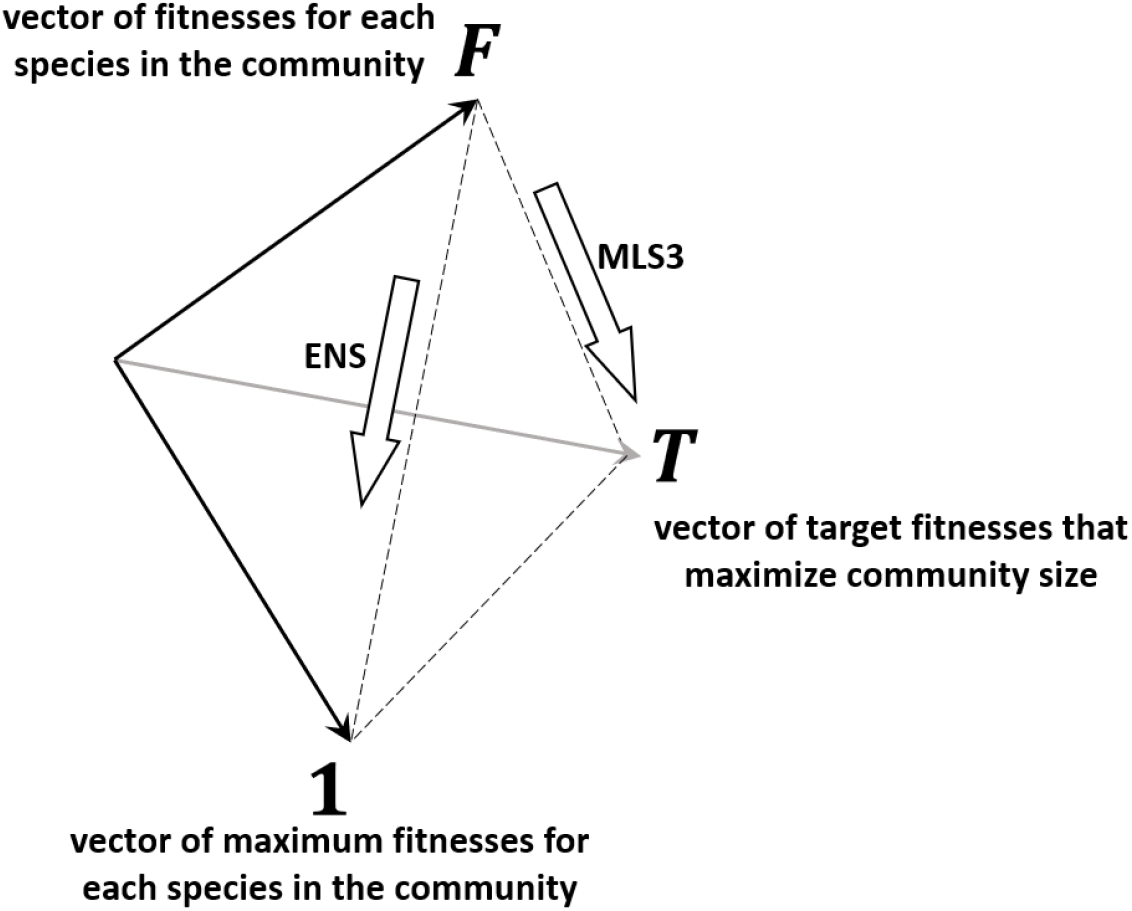
The relationship between the vector ***F*** of fitnesses for each species in a community, the vector ***T*** of target fitnesses that represents the configuration that optimizes community size, and the ones vector **1** that represents the maximum fitnesses for each species. Evolution by natural selection (ENS) acting on individuals would, if unopposed, cause each species to increase in fitness toward its maximum. This can only result in a poorly regulated community with ***F*** = **1**, maximally distant from ***T*** and corresponding to the minimum community size. In a metacommunity setting MLS3 can oppose ENS at the level of individuals or alleles within species by favoring the dispersal of larger communities or those that are better regulated with ***F*** closer to ***T***. ENS and MLS3 are aligned when ***T*** = **1**, and so the ability of MLS3 to promote well-regulated communities is tested only when ***T*** < **1** (i.e., some or all *T_i_* < 1). Note that the tension between ENS and MLS3, represented by the distance between ***T*** and **1**, is absent under both MLS1 and MLS2 due to the assumption that new communities are formed in vacant patches (but only if we can assumed, as we do here, that the timespan is too short for mutation-selection within a community to impact its ***F*** vector).

Simulations were conducted on a 20-by-20 network of ecological patches. Patches in the interior of the network were each connected to their eight nearest neighbors. Edges patches and corner patches had five and three nearest neighbors, respectively. Each community was assumed to interact via colonization or migration with communities occupying neighboring patches only (i.e., spatial relations were explicit, 8). Each patch was initially seeded with a randomly assembled community with each 1 − δ ≤ *F*_*i*_ ≤ 1 drawn from a uniform distribution. This produced a cloud of 400 ***F*** vectors some distance from ***T***. Under what we call “localized” MLS1, patches are visited in random order at each timestep when each community has a probability *P*_*cull*_ = exp (−*N*(*t*)/(5 × 10^5^)) of being culled that is greater for smaller communities. Whenever a cull occurs a member of each species is randomly drawn from among neighboring communities with probabilities proportional to the sizes of those communities. This is equivalent to pooling all neighboring communities and drawing a member of each species in proportion to its number in that pool. One variant of each of the 20 species is selected in this way since a community with less than all 20 species is assumed to be inviable. Localized MLS1 therefore results in a new combination of 20 species variants in the culled patch corresponding to a new ***F*** vector. Larger neighboring communities are more likely to contribute a species to the new community, and so overtime this process causes the ***F*** vectors in the metacommunity to move toward ***T***. Under MLS2, communities are culled in the same manner, but in this case an empty patch is colonized by a replicate of a single neighboring community selected at random with probabilities proportional to the sizes of those communities. A notable limitation of this implementation of MLS2 is that it cannot generate new combinations of species variants. At best it can only select from among the initial set of ***F*** vectors the one closest to ***T***.

Under MLS3 patches are also visited in random order at each timestep, but in this case instead of a chance of being a colonizer there is a probability *P*_*mim*_ = 1 − exp (−*N*(*t*)/(5 × 10^5^)) that the community will “donate” a migration propagule to a randomly selected neighboring patch. Larger communities are more likely to be donors than smaller communities. The migration propagule is assumed to contain cells from all species in the donor community as in MLS2. However, the size of each migrant species population is assumed to be 10% of its population in the donor community. The donor community is therefore replicated in the recipient patch, but at a reduced size. The result is a patch that contains two variants of each species that typically differ in fitness. It is assumed that one variant of each species will eventually be fixed in the recipient patch by a combination of selection and drift. We further assume that this occurs independently for each pair of species variants, and that any adjustment in community size due to changes in ***F*** are realized only after the conflict between each pair has been resolved. Under these simplifications the probability *P*_*mix*_(*S, N*_*m*_, *N*_*m*_) that a donor variant will eventually displace its conspecific counterpart in the recipient patch can be modelled using the diffusion approximation (27) applied to each donor species independently.

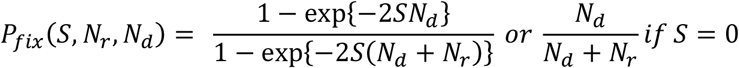

The probability that a donor variant is fixed is a function of both its selection coefficient *S* (the fitness of the donor variant minus the fitness of its conspecific counterpart in the recipient patch) and the proportion *N*_*m*_/(*N*_*m*_ + *N*_*m*_) of cells of the donor type (or mixing ratio, 14), where *N*_*d*_ is the number of donor cells and *N*_*r*_ the number of recipient cells of the same species. Note that *N*_*m*_ + *N*_*m*_ is the effective size of the combined population of donor and recipient genotypes of the same species. Conflict can arise when the donor is less fit that its counterpart (*S* < 0) but originated from a larger community. In such cases the donor might displace its counterpart based on the size of the donor community as reflected by *N*_*m*_. This can cause the ***F*** vectors in the metacommunity to move toward ***T*** and away from the ones vector **1** = ⟨1, …,1⟩ of maximum fitnesses for all 20 species.

To illustrate, suppose a well-regulated community of size 20 × 10^5^ donates 10% of the cells from one of its species populations to a less well-regulated neighboring community of size 20 × 10^4^ cells, so that *N*_*m*_ = *N*_*m*_ = 1 × 10^4^. If the selection coefficient for the donor variant of that species is *S* = −5 × 10^−6^ (i.e., the minimum under our model) then the probability that the donor variant is fixed is *P*_*mix*_(*S, N*_*m*_, *N*_*m*_) = 0.475. Let us now ask, what would the selection coefficient *S*^′^ of a mutant that arises in a single cell in a population of the same size *N* = *N*_*m*_ + *N*_*m*_ have to be to have the same probability of fixation?

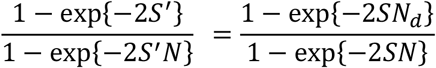

Solving this equation yields *S*^′^ = 0.322 ≫ *S* for the hypothetical mutant. Hence, the size of a migration propagule effectively boosts the selection coefficient of the donor variant if it were to be construed as a mutation in a single cell. The size of the boost diminishes as communities become larger. If *N*_*m*_ = *N*_*m*_ = 1 × 10^5^ for example, the probability that the donor variant is fixed is *P*_*mix*_(*S, N*_*m*_, *N*_*m*_) = 0.270 and the equivalent selection coefficient is *S*^′^ = 0.157. Hence, as the ***F*** vectors in the metacommunity approach ***T*** and the effective population size *N* = *N*_*m*_ + *N*_*m*_ following any migration event increases, the ability of MLS3 to improve community size via the fixation of less fit genotypes is reduced. The maximum mean population size attained by the metacommunity is reached at the point where the effect of selection acting on the fitness of individual species and the opposing effect of the boost in the probability of fixation of less fit variants due to the size *N*_*d*_ of their migrant populations reach an equilibrium. At this point the mean ***F*** vector can no longer move progressively toward the target. The tension between the two levels of selection therefore makes community self-regulation under our model of MLS3 potentially less effective compared to that under our models of MLS1 and MLS2 where that tension is absent.

Six sets of simulations were conducted, each with a different combination of parameter settings (Table 1). In all cases the target vector was constructed as follows:

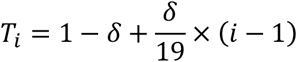

Hence, maximum community size could be reached when *T*_1_ = 1 − δ, *T*_2_ = 1 − δ + δ/19, etc. up to *T*_20_ = 1. Notice that larger values of δ move the targets *T*_*i*_, *i* = 1, …,19 further from the maximum fitness of 1. This was expected to make it more difficult for MLS3 to move ***F*** vectors toward ***T*** because such movement comes at a higher cost in the fitness of members of species 1 to 19. Larger values of *ρ* diminish the strength of the negative correlation between *Dist*(***T, F***(*t*)) and community size by increasing the amplitude of multiplicative noise impacting community size. MLS3 was therefore also expected to be less effective when *ρ* was set to its larger value.

Traditional models of multilevel selection (i.e., MLS1 and MLS2) typically encompass two timescales. In Sober’s (11) verbal model of MLS2, for example, any single population composed of a mix of altruists and selfish types will eventually be composed of the selfish type only. The time interval between colonization events (where populations with more altruists are more likely to be reproduced in vacant patches) must therefore be short compared to the time required for the selfish type to be fixed. The evolutionary dynamic within a patch is consequently assumed to occur over a longer timescale than between-patch dynamics, and all populations are assumed to remain in a state of disequilibrium (i.e., with both altruists and selfish types) until the selfish type goes extinct. We chose to base our model of MLS3 on the diffusion approximation (27) because of its conceptual simplicity, as exemplified by Figure 2. However, our approach also requires the assumption that a population of conspecific organisms never contains more than two genotypes at a time (e.g., a novel type introduced by migration and a wild type already in residence). This means that any novel type introduced into a population must always be fixed or eliminated before the next novel type arrives, and that all communities must therefore have reached equilibrium (i.e., with one variant of each species) before they donate or receive migrants. Our model consequently includes a timescale over which fixation following any migration event occurs, and a potentially longer timescale between migration events. The number of generations before a donor variant is either fixed or eliminated is difficult to estimate. Theorical calculations indicate that the number is bounded above by something on the order of magnitude of the effective population size (28), which in our model can be as much as *N* = 1.1 × 10^6^. Hence, although our results demonstrate how a kind of inter-species altruism can arise in a metacommunity with migration (i.e., under MLS3), the extent to which the time between migration events entailed by our use of the diffusion approximation applies to dispersal in natural metacommunities is not well understood.

The assumptions required by the diffusion approximation can be relaxed by developing other methods for modelling intracommunity dynamics. One approach would be to modify the diffusion approximation to permit migration before fixation. In a population containing two genotypes, for example, the diffusion approximation can be used to calculate the probability distribution *P*(*S, N*_1_(*t*), *N*_2_(*t*), Δ*t*) of the relative numbers of each type after some short time Δ*t* in generations. Numbers *N*_1_(*t* + Δ*t*) and *N*_2_(*t* + Δ*t*) can then be draw from that distribution, representing one timestep. If on the next timestep there is a migration event that donates some number of each type to the population, then the number of each type can be adjusted by summation. New values for *N*_1_ and *N*_2_ can then be drawn from the updated distribution and the process repeated. This approach would still be limited to no more than two variants of any given species at a time but would allow migration to occur over micro-evolutionary timescales as compared to fixation (i.e., since migration could occur before fixation). More flexible alternatives include the use of current models based on single-community dynamics, such as those that use an agent-based approach (e.g., 18, 20, 29) or that are based on some form of Lotka–Volterra equations (e.g., 19, 30). Both approaches can be modified to account for migration between communities in disequilibrium (e.g., 22). Such models would similarly permit migration to occur on a shorter timescale than fixation but have the added advantage that they can also account for more complex intracommunity dynamics (e.g., coexistence of multiple species in the same niche, competitive exclusion). Investigation of these alternatives is left for future efforts.

## FUNDING

This work was supported by grant NFRFE-2019-00703 from the New Frontiers Research Fund of Canada.

## Notes

### Competing Interest Statement

The authors have declared no competing interest.

